# Lessons learned from bugs in models of human history

**DOI:** 10.1101/2020.06.04.131284

**Authors:** Aaron P. Ragsdale, Dominic Nelson, Simon Gravel, Jerome Kelleher

## Abstract

Simulation plays a central role in population genomics studies. Recent years have seen rapid improvements in software efficiency that make it possible to simulate large genomic regions for many individuals sampled from large numbers of populations. As the complexity of the demographic models we study grows, however, there is an ever-increasing opportunity to introduce bugs in their implementation. Here we describe two errors made in defining population genetic models using the msprime coalescent simulator that have found their way into the published record. We discuss how these errors have affected downstream analyses and give recommendations for software developers and users to reduce the risk of such errors.

## Introduction

In the effort to build more realistic simulations of genetic diversity, scientific software developers often focus on computational speed and biological realism. As the models simulated become more realistic, however, they also become more complex and difficult to specify. The interface through which users define their models is therefore increasingly important. Without an intuitive and thoroughly documented interface, it is very difficult to simulate complex population models correctly.

The `msprime` coalescent simulator (Kelleher et al., 2016; Nelson et al., 2020; Kelleher and Lohse, 2020) is now widely used in genetics studies. Much of its appeal is the large increase in efficiency over the classical `ms` program (Hudson, 2002) which makes it feasible to simulate large samples of whole chromosomes for the first time. Another distinct advantage of `msprime` is its Python application programming interface (API), which greatly increases the flexibility and ease of use over the standard approach of text-based command line interfaces. In particular, programs like `ms` require users to specify cryptic command line options to describe demographic models. For example, the Gutenkunst et al. (2009) demographic model (that is the subject of this note), as written in `ms` syntax, is

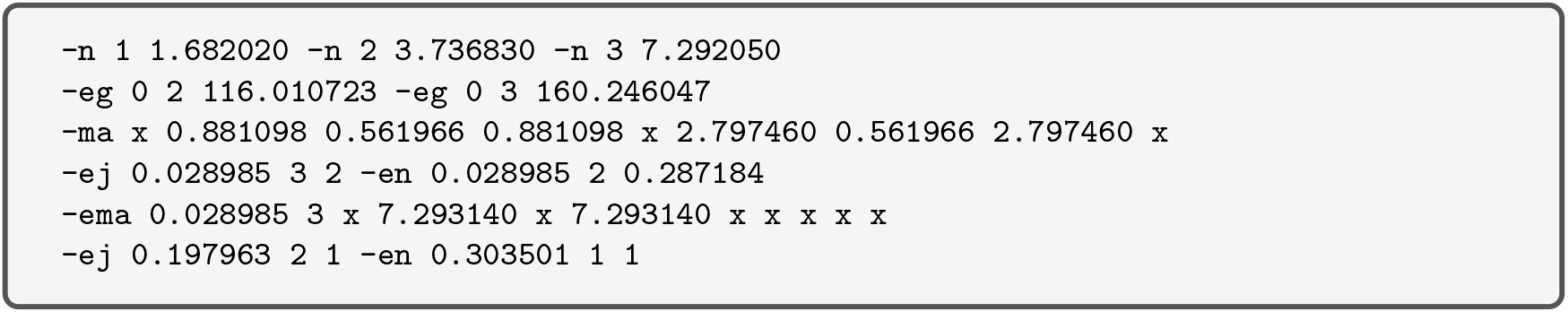

This model is relatively simple, and models with many more populations and parameters are increasingly common. Descriptions of such models in `ms` syntax are not easy to comprehend. The Python interface for `msprime`, by contrast, allows the user to state models in a more human-readable and programmatic manner, and has many advantages over `ms`’s command line interface.

Even when using a high-level programming language like Python, however, implementing multi-population models of demographic history is difficult and error-prone. In this note we discuss two implementation errors that arose through unfortunate design decisions in `msprime`’s demography API and which then found their way into the scientific record. The first error has relatively mild effects on genetic diversity but was used in many publications, while the second error was used only once but had a large impact on the simulation results. In light of these implementation errors, we discuss improvements to `msprime`’s API motivated by these discoveries and, more generally, best practices for implementing and simulating complex multi-population demography.

## Case 1: A misspecified model in msprime’s documentation

To illustrate the demography API, `msprime` included a description of a widely-used three population Out-of-Africa model (Gutenkunst et al., 2009) as part of its tutorial documentation. In this model (Fig. 1A), Eurasian (CEU and CHB) and African (YRI) populations split from each other in the deep past, followed by a more recent split of European and Asian populations, with variable rates of continuous migration between each of the populations. Regrettably, the implementation in the `msprime` tutorial was incorrect. Before the time of the split of African and Eurasian populations, when there should have been just a single randomly mating population, migration was allowed to occur between the ancestral population and a second population with size equal to the Eurasian bottleneck size for all time into the past (Fig. 1B). This incorrect model was introduced into the tutorial for `msprime` ver. 0.3.0 and remained in the documentation for around four years.

**Figure 1:**
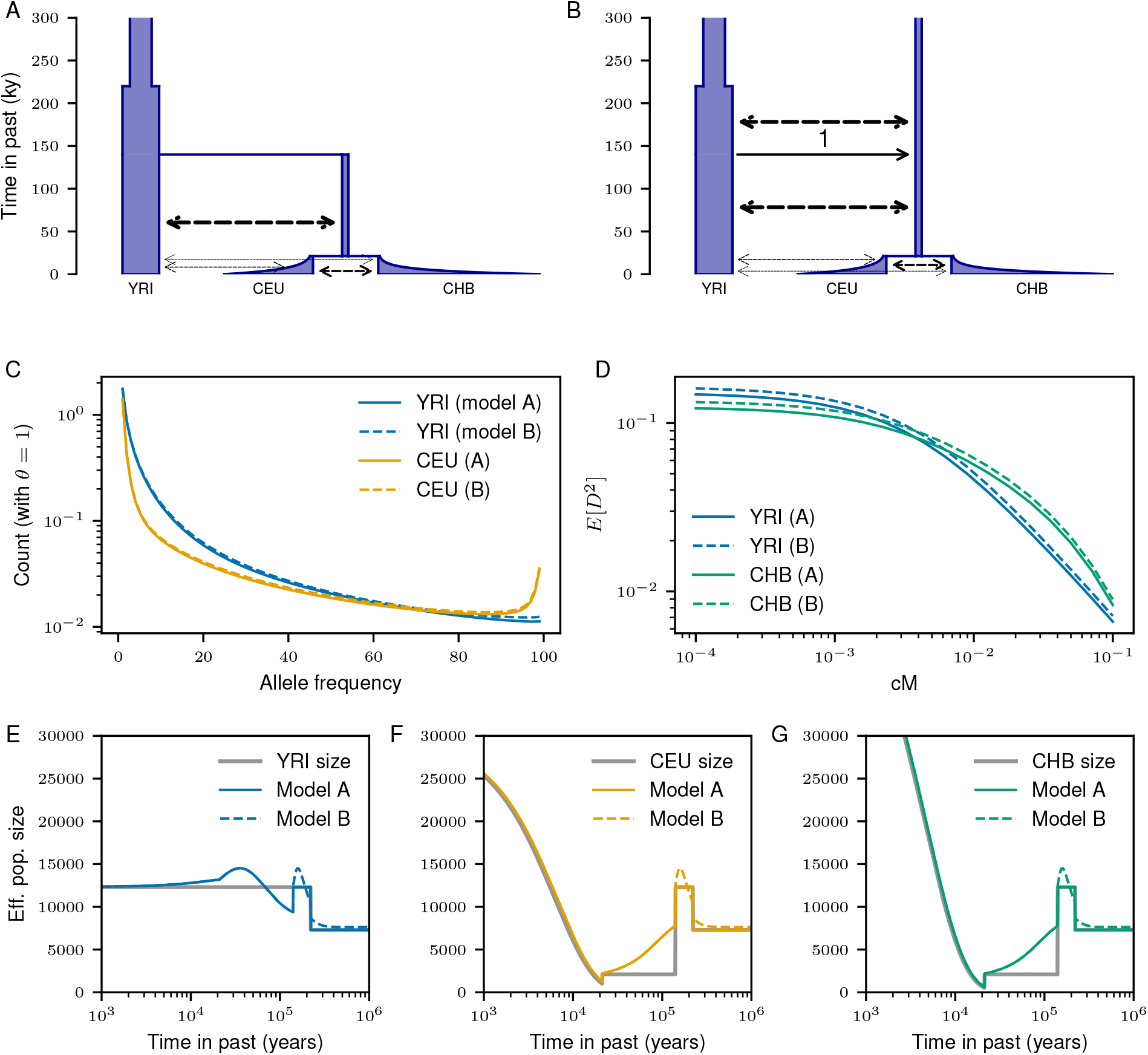
Expected diversity statistics under the Gutenkunst et al. (2009) model. (A) The correctly implemented model. Dashed arrows depict continuous migration. (B) The incorrectly implemented model from the `msprime` tutorial, with migration continuing into the past beyond the mass migration event with proportion 1 from the ancestral population to the bottleneck population. (C) Marginal allele frequency spectra under the two models. Heterozygosity in the incorrect model is inflated by ~ 3. 5%, though the general shape of the distributions are qualitatively similar. (D) Similarly, the increased heterozygosity leads to excess *D*^2^, though the LD-decay is qualitatively similar between models. (E-G) True size history for each population plotted against the expected size history computed from the inverse coalescence rates under each model.

Fortunately, the effects of this error are subtle. Population sizes and structure since the time of the earliest split are unaffected, so differences in expected *F_ST_* are negligible between the correct and incorrect models. However, the ancient structure distorts the distribution of early TMRCAs (Fig. 1E-G). The extraneous ancient population increases the long-term effective population size, resulting in roughly 4% excess heterozygosity in contemporary populations compared to the intended model, but the overall effects on patterns of diversity are minimal (Fig. 1C,D).

Even though the error has a limited effect on simulated data, the tutorial code has been copied many times and used in publications. By searching for some identifying strings from the model definition on GitHub, we found 32 repositories containing either direct copies of the erroneous model code, or code that was obviously derived from it. (We have opened issues on each of these repositories to alert the authors.) In most cases the publications used simulations to test a non-demographic inference method, and the model was used as an example of a complex population history (Kelleher et al., 2019; Albers and McVean, 2020; Tong and Hernandez, 2020). Zhou et al. (2018) used the incorrect model as an example of how their method for visualising demographic models can support `msprime` input. Finally, Pfaffelhuber et al. (2020) used simulations of the incorrect model demography to evaluate their method for choosing ancestry informative markers. Given the very subtle effect of the incorrect model on demography (and the fact the method was evaluated using other simulations and real data), it seems unlikely that the model details had any qualitative effect on their conclusions.

This long-standing error could have been prevented by better API design. To model a population split currently in `msprime`, a user must specify a `MassMigration` event that moves lineages from one population to another and then must also remember to turn off migration between those populations at the same time. The demographic events for the correct model are given as

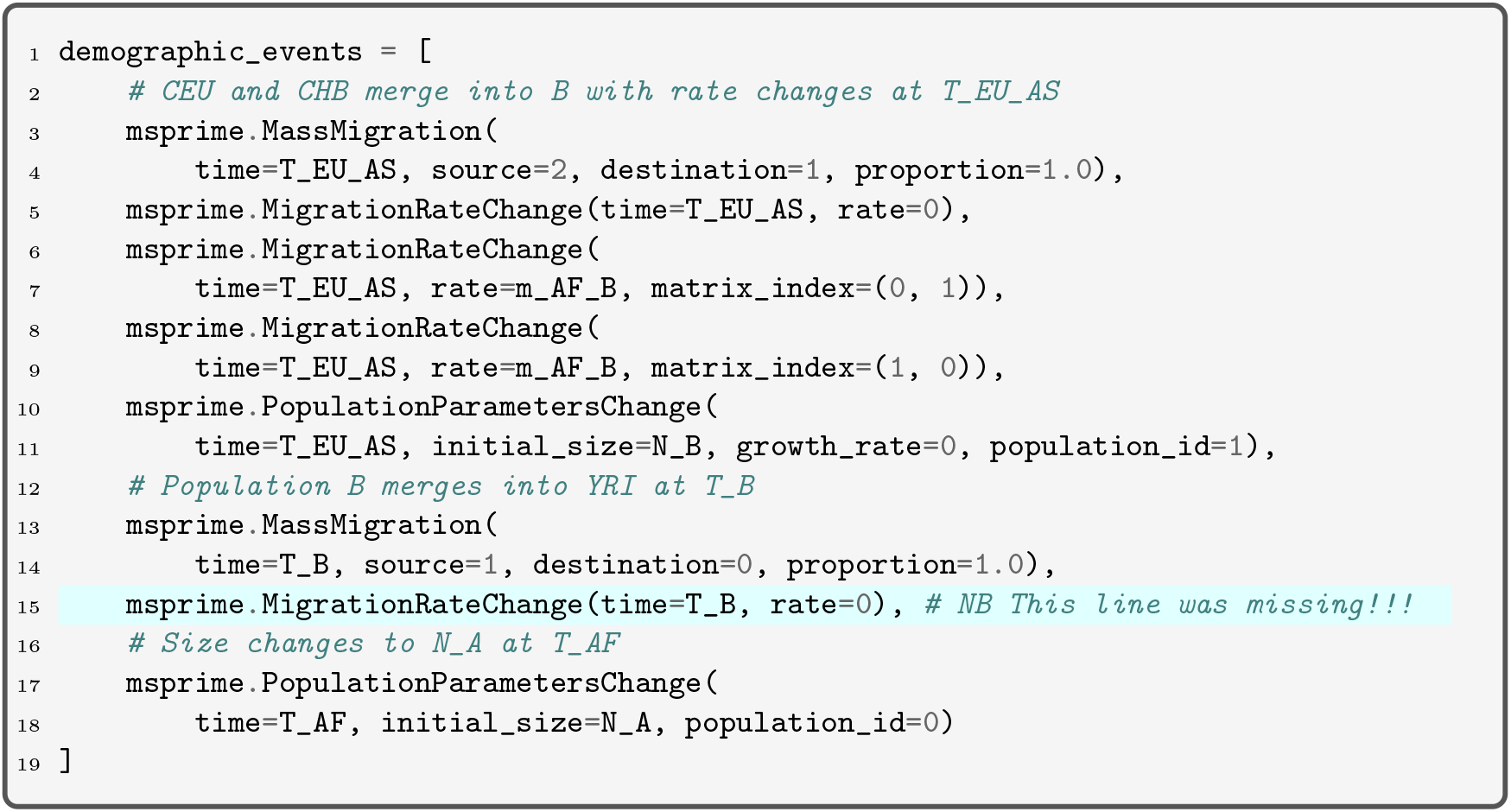

Note that the migration rate change on 15 was missing in the incorrect model. The release of `msprime` version 1.0 will introduce a `PopulationSplit` event, which more intuitively links the movement of lineages with appropriate changes in migration rates at the time of the split.

### Case 2: Incorrect model parameters in an analysis pipeline

In another publication using this model (Martin et al., 2017), a separate error was introduced: the model itself was defined as suggested in the documentation (using updated parameters from Gravel et al. (2011)) and inspected using the `msprime` debugging tools. Despite these initial checks being made, the simulation was performed without passing the parameter defining demographic events, so that the three populations never merged and remained separated with low levels of migration (Fig 2A), leading to a vast overestimate of the divergence across human populations. While the correct model predicts a mean *F_ST_* of 0.05-0.10 across the three population, the simulated model generated *F_ST_* ranging between 0.3-0.6, depending on the populations considered. Overall diversity was also strongly affected: heterozygosity was more than doubled in African populations and reduced by more than half in Eurasian populations relative to the correct model.

**Figure 2:**
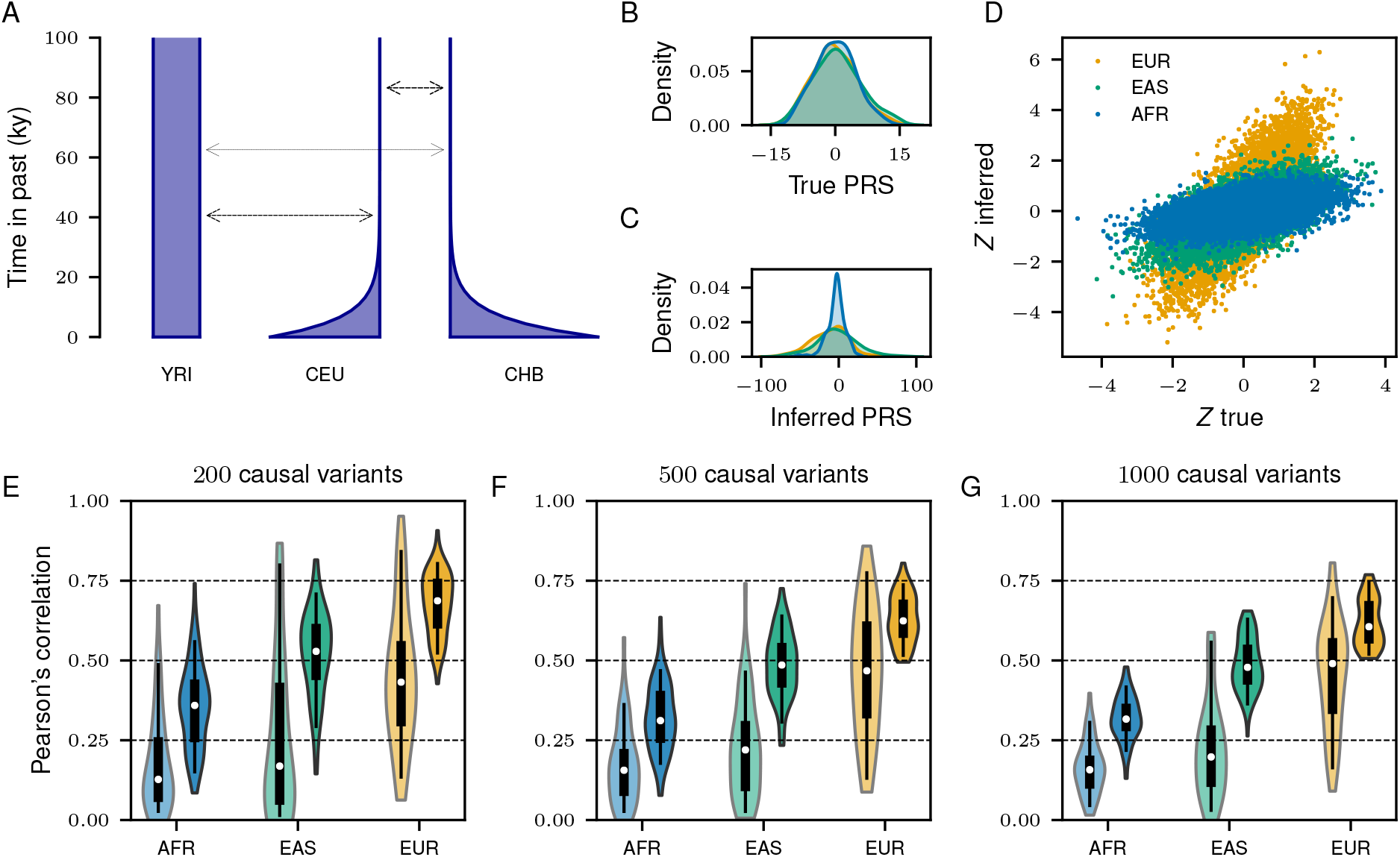
The transferability of PRS under neutrality. **(A)**In Martin et al. (2017), the simulated demographic model did not apply demographic events in the past, so continental populations were simulated as isolated with low levels of migration for all time. The correct model has historical events as shown in Fig. 1A. **(B-G)**We repeated the simulation experiment in Martin et al. (2017) using the correct demographic model. GWAS summary statistics were computed from 10,000 case and control subjects in the European population, and the distribution of inferred PRS were compared across the three simulated populations. Correlations between true and inferred polygenic scores were computed over 100 simulation replicates. Unlike the original study, we do not observe large differences in mean inferred PRS across the three populations (C, D), and while risk prediction in the African and East Asian populations is still reduced compared to the European population, the reduction in prediction accuracy is not as large as reported in the original study (E-G). Panel D shows a comparison of PRS distributions for a simulation replicate with 1,000 causal variants. For each population in the violin plots, the original correlations from Martin et al. (2017) are shown on the left, and correlations using the correct model are shown on the right. For direct comparison to the original study, see Figure 5 in Martin et al. (2017).

This simulation was performed to assess the transferability of polygenic risk scores across human populations. In other words, it sought to explore how human demographic history and population structure affect our ability to predict genetic risk in diverse populations given the well-documented unequal representation in medical genetic studies (Popejoy and Fullerton, 2016). The resulting publication has been influential in the discussion of health inequalities and genomics, with over 350 citations since 2017. The large excess in divergence under the incorrect model here exaggerated the role of demography and genetic drift in limiting the transferability of genetic risk scores across populations.

Difficulties in transferability remain in the corrected model (Fig. 2), although risk prediction in each population is significantly improved (compare to Fig. 5 in Martin et al. (2017)). Simulations under the correct model indicate that the accuracy of genetic risk scores is still substantially reduced in understudied populations (Fig. 2E-G), supporting one of the main conclusions of Martin et al. (2017). However, the reduction is much less pronounced than reported. In particular, we do not observe large differences in mean predicted risk across populations (Fig. 2C,D), present in data and simulations from Martin et al. (2017). Thus, a model with continental-scale demographic structure, highly polygenic architecture, and neutral evolution does not appear to explain the large directional biases in mean predicted risk identified in Martin et al. (2017).

From a software perspective, this error was easy to make and could have been prevented by better API design. The original `msprime` API requires the user to pass three separate parameters to specify a demographic model. For example,

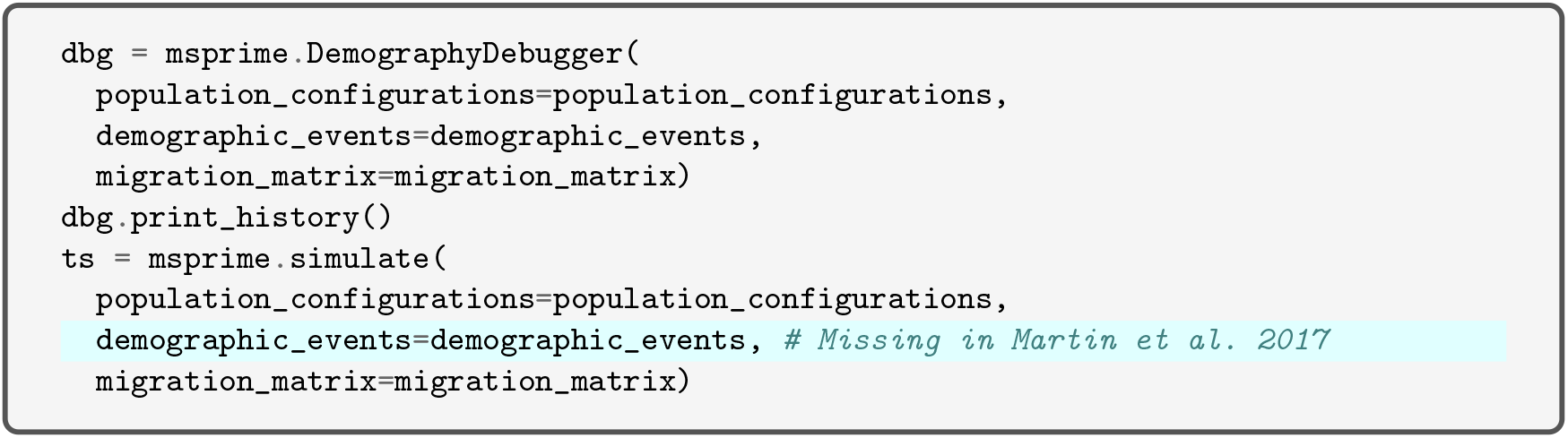

Note that the same three parameters must be passed to both the debugger and the simulate function. To help prevent such errors, `msprime` ver. 1.0 will introduce a Demography class. The above snippet would then be rewritten as

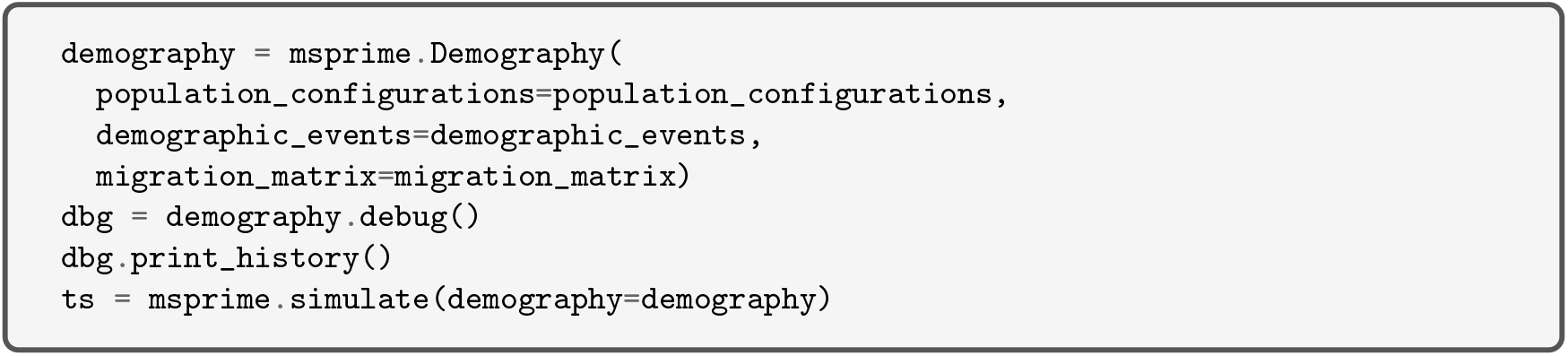

This simple change to the interface makes it much less likely that different models are passed to the debugger and simulator, reducing the potential for error.

## Conclusions

The implementation of complex demographic models is error prone, and such errors can have a large impact on downstream analyses and interpretation. The discovery and correction of the demographic models discussed here underscore how API design choice can lead to the propagation of mistakes that are difficult to notice. We therefore recommend the following steps to ensure more robust simulations.

Firstly, if possible, we recommend avoiding the implementation of demographic models entirely by using `stdpopsim` (Adrion et al., 2019). This is a “standard library” of quality-controlled models and simulation resources for a growing number of commonly studied species. The resource is built around an open-source community development model with rigorous code review and quality control procedures. For example, when a contributor adds a new model it is not fully integrated into the catalog until a second, entirely independent, implementation of the model is provided by another developer. If the original and “QC” versions of the model are precisely equal, then we can have a reasonable degree of confidence in correctness. It is through this QC process that we discovered the misspecified model in `msprime`’s documentation. If the required model is not present in `stdpopsim`, then particular care should be taken to validate the implementation. This could include verification through code review or comparison with an independent implementation of the model. If the model is of wider interest to the community, we would encourage developers to contribute it to `stdpopsim`: at the very least, the quality-control procedures in place provide a strong reassurance of correctness.

Secondly, regardless of whether the model has been implemented locally or obtained from a resource like `stdpopsim`, basic statistical validation should *always* be performed on the results. Errors are all too easy to make, and an analysis of the basic statistical properties of the simulations is essential due diligence. For example, a brief analysis of *F_ST_* values would likely would have prevented the error in Martin et al. (2017). It is understandable that such analyses were not undertaken in this case, as processing such a huge dataset (600,000 samples) was highly challenging. Recent progress has made statistical analysis this scale much easier (Ralph et al., 2020), but the problems would have been apparent if smaller pilot simulations were undertaken and analysed. Graphical inspection of demographic models can also help identify issues, especially if visualization can be automated. The demography debugging tool in `msprime` summarizes demographic events occurring over time, and various efforts are underway to facilitate the specification and visualization of population genetics models, such as Zhou et al. (2018) and the demography package used in this paper.

Finally, openness is essential to the self-correcting nature of science. We only know about these errors because of open code and open-source development processes. By making their entire pipeline available, Martin et al. (2017) not only enabled other research teams to build upon their findings, but they made it possible for errors to be found and corrected. There must be many, many more mistakes out there, and we need both pre- and post-publication vigilance from users and developers to ensure the soundness of the large body of simulation-based analyses.

## Methods

We computed the expected allele frequency spectrum using `moments` ver. 1.0.3 (Jouganous et al., 2017), and LD-decay curves using `moments.LD` (Ragsdale and Gravel, 2019). *F_ST_* and other diversity statistics were computed from the expected AFS and verified using branch statistics from the output of `msprime` simulations using tskit ver. 0.2.3 (Ralph et al., 2020). Demographic models were plotted using the demography package (https://github.com/apragsdale/demography, ver.0.0.3). We used the original pipeline from Martin et al. (2017) available from https://github.com/armartin/ancestry_pipeline/blob/master/simulate_prs.py. We updated the pipeline to run with the correct demographic parameters and more recent versions of `msprime` and tskit, and the updated pipeline is available at https://github.com/apragsdale/PRS. Data and Python scripts to recreate Figures 1 and 2 can be found at https://github.com/jeromekelleher/msprime-model-errors. A full list of the GitHub repositories containing copies of the erroneous model are also given here.

## Acknowledgements

We would like to thank Alicia Martin for comments on the manuscript and for providing the simulation data from Martin et al. (2017) used in Fig. 2. We also like to thank Chris Gignoux and Ilan Gronau for helpful comments on the manuscript. JK would like to apologise to those affected by the error in the example Out-of-Africa model and by the poor design decisions in `msprime`’s demography API.

## References

Jeffrey R Adrion, Christopher B Cole, Noah Dukler, Jared G Galloway, Ariella L Gladstein, Graham Gower, Christopher C Kyriazis, Aaron P Ragsdale, Georgia Tsambos, Franz Baumdicker, et al. A community-maintained standard library of population genetic models. bioRxiv, 2019.

Patrick K Albers and Gil McVean. Dating genomic variants and shared ancestry in population-scale sequencing data. PLoS biology, 18(1):e3000586, 2020.

Simon Gravel, Brenna M Henn, Ryan N Gutenkunst, Amit R Indap, Gabor T Marth, Andrew G Clark, Fuli Yu, Richard A Gibbs, Carlos D Bustamante, 1000 Genomes Project, et al. Demographic history and rare allele sharing among human populations. Proceedings of the National Academy of Sciences, 108(29):11983–11988, 2011.

Ryan N Gutenkunst, Ryan D Hernandez, Scott H Williamson, and Carlos D Bustamante. Inferring the joint demographic history of multiple populations from multidimensional snp frequency data. PLoS genetics, 5(10):e1000695, 2009.

Richard R. Hudson. Generating samples under a Wright-Fisher neutral model of genetic variation. Bioinformatics, 18(2):337–338, 2002.

Julien Jouganous, Will Long, Aaron P Ragsdale, and Simon Gravel. Inferring the joint demographic history of multiple populations: beyond the diffusion approximation. Genetics, 206(3):1549–1567, 2017.

Jerome Kelleher and Konrad Lohse. Coalescent simulation with msprime. In Julien Y. Dutheil, editor, Statistical Population Genomics, pages 191–230. Springer US, New York, NY, 2020.

Jerome Kelleher, Alison M Etheridge, and Gilean McVean. Efficient coalescent simulation and genealogical analysis for large sample sizes. PLoS computational biology, 12(5):e1004842, 2016.

Jerome Kelleher, Yan Wong, Anthony W Wohns, Chaimaa Fadil, Patrick K Albers, and Gil McVean. Inferring whole-genome histories in large population datasets. Nature genetics, 51(9):1330–1338, 2019.

Alicia R Martin, Christopher R Gignoux, Raymond K Walters, Genevieve L Wojcik, Benjamin M Neale, Simon Gravel, Mark J Daly, Carlos D Bustamante, and Eimear E Kenny. Human demographic history impacts genetic risk prediction across diverse populations. The American Journal of Human Genetics, 100(4):635–649, 2017.

Dominic Nelson, Jerome Kelleher, Aaron P Ragsdale, Claudia Moreau, Gil McVean, and Simon Gravel. Accounting for long-range correlations in genome-wide simulations of large cohorts. PLoS genetics, 16 (5):e1008619, 2020.

Peter Pfaffelhuber, Franziska Grundner-Culemann, Veronika Lipphardt, and Franz Baumdicker. How to choose sets of ancestry informative markers: A supervised feature selection approach. Forensic Science International: Genetics, 46:102259, 2020.

Alice B Popejoy and Stephanie M Fullerton. Genomics is failing on diversity. Nature News, 538(7624): 161, 2016.

Aaron P Ragsdale and Simon Gravel. Models of archaic admixture and recent history from two-locus statistics. PLoS genetics, 15(6):e1008204, 2019.

Peter Ralph, Kevin Thornton, and Jerome Kelleher. Efficiently summarizing relationships in large samples: a general duality between statistics of genealogies and genomes. Genetics, 2020.

Dominic MH Tong and Ryan D Hernandez. Population genetic simulation study of power in association testing across genetic architectures and study designs. Genetic epidemiology, 44(1):90–103, 2020.

Ying Zhou, Xiaowen Tian, Brian L Browning, and Sharon R Browning. POPdemog: visualizing population demographic history from simulation scripts. Bioinformatics, 34(16):2854–2855, 2018.

